# Cardiomyocyte alpha-1A adrenergic receptors mitigate post-infarct remodeling and mortality by constraining necroptosis

**DOI:** 10.1101/2022.08.29.505687

**Authors:** Jiandong Zhang, Peyton B. Sandroni, Wei Huang, Leah Oswalt, Alan J. Smith, Tyler Ash, Sung-Hoo Lee, Yen-Yu I. Shih, Joseph S. Rossi, Hsiao-Ying S. Huang, Bat E. Myagmar, Paul C. Simpson, Jonathan C. Schisler, Brian C. Jensen

**Affiliations:** University of North Carolina School of Medicine, Division of Cardiology; UNC McAllister Heart Institute; Mechanical and Aerospace Engineering Department, North Carolina State University; Center for Animal MRI, University of North Carolina; San Francisco VA Medical Center; University of California, San Francisco; UNC Department of Pharmacology

**Author notes:** To whom correspondence should be addressed: Brian C. Jensen MD, 160 Dental Circle, CB 7075, Chapel Hill, NC 27599-7075.

## Abstract

Activation of alpha-1-adrenergic receptors (α1-ARs), particularly the α1A subtype, protects the murine heart against injury, whereas human studies show that α1-AR antagonists (α-blockers) may increase the risk of heart failure. We created a cardiomyocyte-specific α1A-AR knockout mouse (cmAKO) to define the mechanisms underlying these effects and to elucidate whether they arise from cardiomyocyte α1A-ARs or systemic factors. Myocardial infarction (MI) resulted in 70% 7-day mortality in cmAKO compared to 10% in wild type (WT) mice. cmAKO mice exhibited exaggerated ventricular remodeling and increased cell death compared to WT mice 3 days post-MI, coupled to upregulation of canonical mediators of necroptosis: receptor-interacting protein (RIP) kinases RIP1 and RIP3 and mixed lineage kinase domain-like protein. An α1A-AR agonist mitigated ischemia-induced cardiomyocyte death and necroptotic signaling *in vitro*. A RIP1 antagonist abrogated the protective effects of α1A activation *in vivo* and *in vitro*. We found that patients at our center who were taking α-blockers at the time of MI experienced a higher risk of mortality (hazard ratio 1.53, p=0.029) during 5-year follow-up, providing clinical correlation for our experimental data. Collectively our findings indicate that cardiomyocyte α1A-ARs constrain ischemia-induced necroptosis and suggest caution in the use of α-blockers in patients at risk for MI.

## Introduction

Despite significant advances in clinical management, acute myocardial infarction (MI) remains a leading source of morbidity and mortality worldwide(1) and the number of patients who develop heart failure (HF) after MI is increasing(2). Acute and subacute cardiomyocyte death are central to the pathobiology of MI and collectively determine the extent of post-infarct remodeling and the likelihood of consequent HF. Historically, cell death in the setting of MI has been attributed largely to passive necrosis and apoptosis, but there is increasing recognition that programmed necrosis (necroptosis) contributes meaningfully to post-infarct injury and remodeling(3).

Activation of the sympathetic nervous system (SNS) is well-established as a critical component of the physiological response to MI in both experimental animal models and humans(4–6). SNS activation is mediated by increased levels of the catecholamines norepinephrine (NE) and epinephrine (EPI) that regulate their critical functions through binding to two major categories of adrenergic receptors (ARs) in the heart, alpha-1-ARs (α1-ARs) and beta-AR (β-ARs). Sustained hyperactivation of β1-ARs is a toxic driving force contributing to excessive cardiomyocyte death after MI and β-blockers have long been recognized as cornerstones of medical therapy in the immediate post-MI period(7). The potential roles of cardiomyocyte α1-ARsin this context have received considerably less attention, but a growing body of experimental data indicates that α1-ARs protect against myocardial ischemic injury (reviewed in (8, 9)).

There are three subtypes of α1-ARs with distinct tissue distribution and physiological function in the heart. The α1A and α1B are the predominant α1-AR subtype in rodent and human myocardium(10, 11), whereas the α1D is found in coronary arterial smooth muscle(12, 13). Work published in the 1990s demonstrated that non-selective α1-AR activation conferred protection against ischemia-reperfusion injury and limited post-infarct remodeling(14, 15). Subsequent studies identified the α1A as the cardioprotective α1-AR subtype in the context of both ischemic and non-ischemic insults (reviewed in (16)). Overexpression of the α1A in mice(17) and rats(18) limits infarct size and protects contractile function after MI. Global α1A knockout mice exhibit exacerbated post-infarct remodeling and apoptosis four weeks after left coronary artery (LCA) ligation(19), but the response to injury has not been studied in mice with cardiomyocyte-specific α1A deletion.

Here we have used Cre-lox breeding to develop a cardiomyocyte-specific α1A knockout (cmAKO) mouse line, obviating valid concerns about potential physiological confounding in global α1-AR loss-of-function models. We find that cmAKO mice experience markedly increased one-week mortality after MI due to permanent LCA ligation, with evidence of exaggerated ventricular remodeling by Day 3. *In vivo* and *in vitro* studies demonstrated enhanced activation of RIP kinase-dependent necroptosis pathways in cmAKO mice, revealing a novel mechanism for α1A-mediated cardioprotection. We also show that patients who were taking medications that block α1A-AR signaling at the time of percutaneous coronary intervention (PCI) for MI experienced a higher risk of mortality during a 5-year follow-up period, providing clinical correlation for our experimental findings.

## Results

### Generation of mice with specific deletion of the α1A-AR in cardiomyocytes

To examine the functions of cardiomyocyte α1A-ARs, we created a cardiomyocyte-specific α1A-AR knockout mouse line (cmAKO). We initially bred mice with Cre recombinase under the control of a truncated *Myh6* promoter(20) with mice carrying the ROSA^mT/mG^ Cre-reporter allele, allowing us to confirm Cre expression (green fluorescent protein GFP+) within the ventricle of *Myh6*-Cre/ROSA^mT/mG^ mice but the absence of Cre expression (red fluorescent protein RFP+) in noncardiac tissues such as brain, kidney and spleen (**Figure 1A**). To generate the cmAKO mice we bred the *Myh6*-Cre mice with a mouse line harboring loxP sites flanking the first coding exon in *Adra1a* (the gene encoding the α1A-AR). To confirm cardiac specific deletion of the α1A-AR in our cmAKO mice, we isolated RNA from heart as well as brain, kidney and spleen. Using quantitative reverse transcriptase PCR (qRT-PCR), we determined that cmAKO mice exhibited a >90% reduction in α1A-AR mRNA in the heart, but preserved α1A-AR expression in all other tissues examined (**Figure 1B**). Expression of the α1B-AR *(Adra1b)* in the left ventricle was unaffected by the Cre-lox recombination event (**Figure 1C**). For all experiments in this manuscript, we used *Myh6-* Cre^neg^/*Adra1a*^flox/flox^ mice as wild type (WT) controls for our *Myh6*-Cre^pos^/*Adra1a*^flox/flox^ mice (cmAKO).

**Figure 1.**
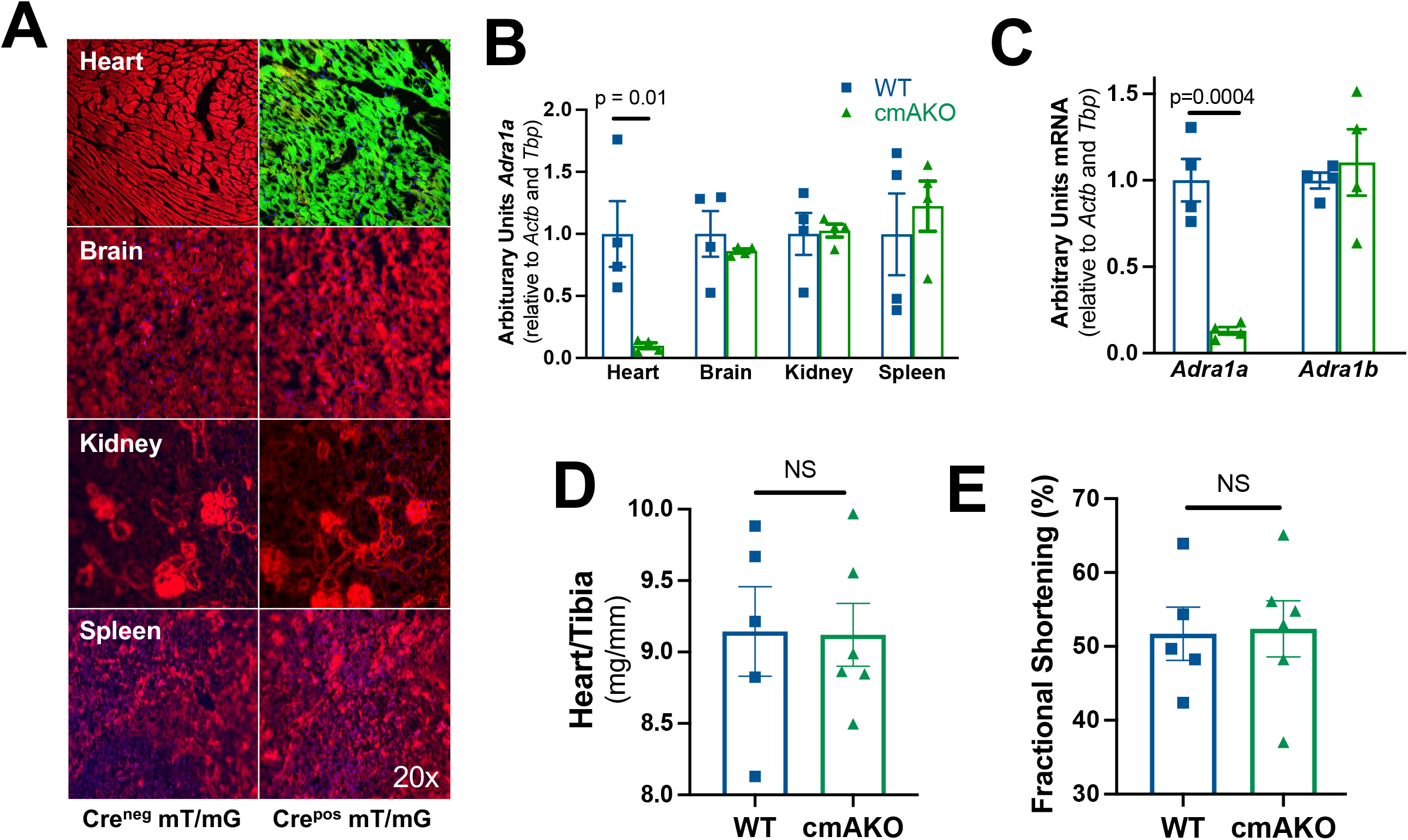
Validation of mice lacking cardiomyocyte α1A-ARs. **(A)** Representative images of heart, brain, and kidney in *Myh6*-Cre/ROSA^mT/mG^ mice. Green fluorescence (GFP) indicates the presence of *Myh6*-Cre expression (Cre^pos^) whereas red fluorescence (RFP) indicates the absence of *Myh6*-cre expression (Cre^neg^). **(B)** mRNA expression for the α1A-AR (*Adra1a*) on tissues from WT and cmAKO littermates; **(C)** mRNA expression of α1A-AR and α1B-AR *(Adra1b)* in left ventricles of WT and cmAKO mice; **(D)** mRNA expression in isolated adult mouse ventricular myocytes; **(E)** Fractional shortening, as measured by conscious echocardiography. *Actb* = beta-actin; cmAKO = cardiomyocyte-specific *Adra1a* knockout mouse; *Tbp* = TATA binding protein; WT = wild type (αMHC-Cre^neg^/α1A-AR^fl/fl^).

### Absence of cardiomyocyte α1A-ARs does not alter baseline cardiac mass or function

Next, we examined the baseline cardiac function of our cmAKO mice and their littermates. As shown in **Table 1**, the absence of cardiomyocyte α1A did not affect either body weight or heart weight. Heart weight indexed to tibia length was identical in cmAKO mice and their littermates (**Figure 1D**), consistent with published phenotypes for global α1A-KO and α1A transgenic mice. Using conscious echocardiography to further characterize the cardiac structure and function of the cmAKO mice, we found that fractional shortening was 52 ± 8% in WT and 52 ± 9% in cmAKO mice (**Table 1**, **Figure 1E**). Collectively these findings strongly suggest that the physiological cardiac hypertrophy induced by non-selective α1-AR agonists(9) is mediated by the α1 B subtype. They also indicate that though enhancing α1A activity preserves contractile function in the setting of injury(8), its presence is not required to maintain normal basal contractile function.

**Table 1.** Basal cardiac parameters in WT and cmAKO mice (n=6 per group)

|  | **WT** | **cmAKO** | **p** |
| --- | --- | --- | --- |
| Heart Weight (mg) | 153 ± 13 | 157 ± 9 | NS |
| Body Weight (g) | 25.8 ± 2.1 | 25.9 ± 1.7 | NS |
| Tibia Length (mm) | 16.7 ± 0.7 | 17.2 ± 0.6 | NS |
| HW/BW (mg/g) | 5.9 ± 0.8 | 6.1 ± 0.4 | NS |
| HW/TL (mg/mm) | 9.1 ± 0.7 | 9.1 ± 0.5 | NS |
| HR (bpm) | 694 ± 37 | 634 ± 77 | NS |
| LV mass (mg) | 113 ± 5 | 103 ± 18 | NS |
| EF (%) | 83.4 ± 6.7 | 83.9 ± 8.5 | NS |
| FS (%) | 51.7 ± 8.0 | 52.4 ± 9.3 | NS |

BW=body weight; EF=ejection fraction by echocardiogram; FS=fraction shortening by echocardiogram; HR=heart rate; HW=heart weight; LV=left ventricle; NS=not significant. All data were expressed as mean ± SEM. An unpaired t-test compared the two groups in each instance.

### Cardiomyocyte α1A-ARs protect against early injury in a permanent left coronary artery ligation model

After verification and basal characterization of our cmAKO and WT mice, we sought to test whether the absence of cardiomyocyte α1A-ARs affected the response to injury in an experimental model of MI. Towards this end, male cmAKO mice and their littermates were subjected to a well-established LCA ligation model. Of 15 WT and 17 cmAKO mice that underwent surgery, only 2 WT mice died between postoperative day 3 and day 10, however 11 cmAKO mice died during this period. At 14 days post-infarction, survival in WT mice was 86.7% whereas survival in cmAKO mice was 35.3% (p=0.005, **Figure 2A**). Given the striking difference in mortality, a subgroup of dead mice was subjected to necropsy with histological examination. We found evidence of left ventricular free wall rupture in 6 of 9 cmAKO mice, as confirmed using published validated criteria(21): coagulated blood in the chest cavity with disruption of the ventricle on gross examination (**Supplemental Figure S1A**) and extensive coagulative necrosis on histopathological assessment (**Supplemental Figure S1B**). In light of the excessive mortality by Day 5, we turned our attention thereafter to an earlier timepoint, conducting all subsequent analyses at Day 3 post-ligation. Using established protocols for measuring infarct area(22), we found that cmAKO mice sustained 50% larger infarcts than WT mice (22 ± 2% LV mass vs 34 ± 3% LV mass, p=0.015) by post-MI Day 3 (**Figure 2B**).

**Figure 2.**
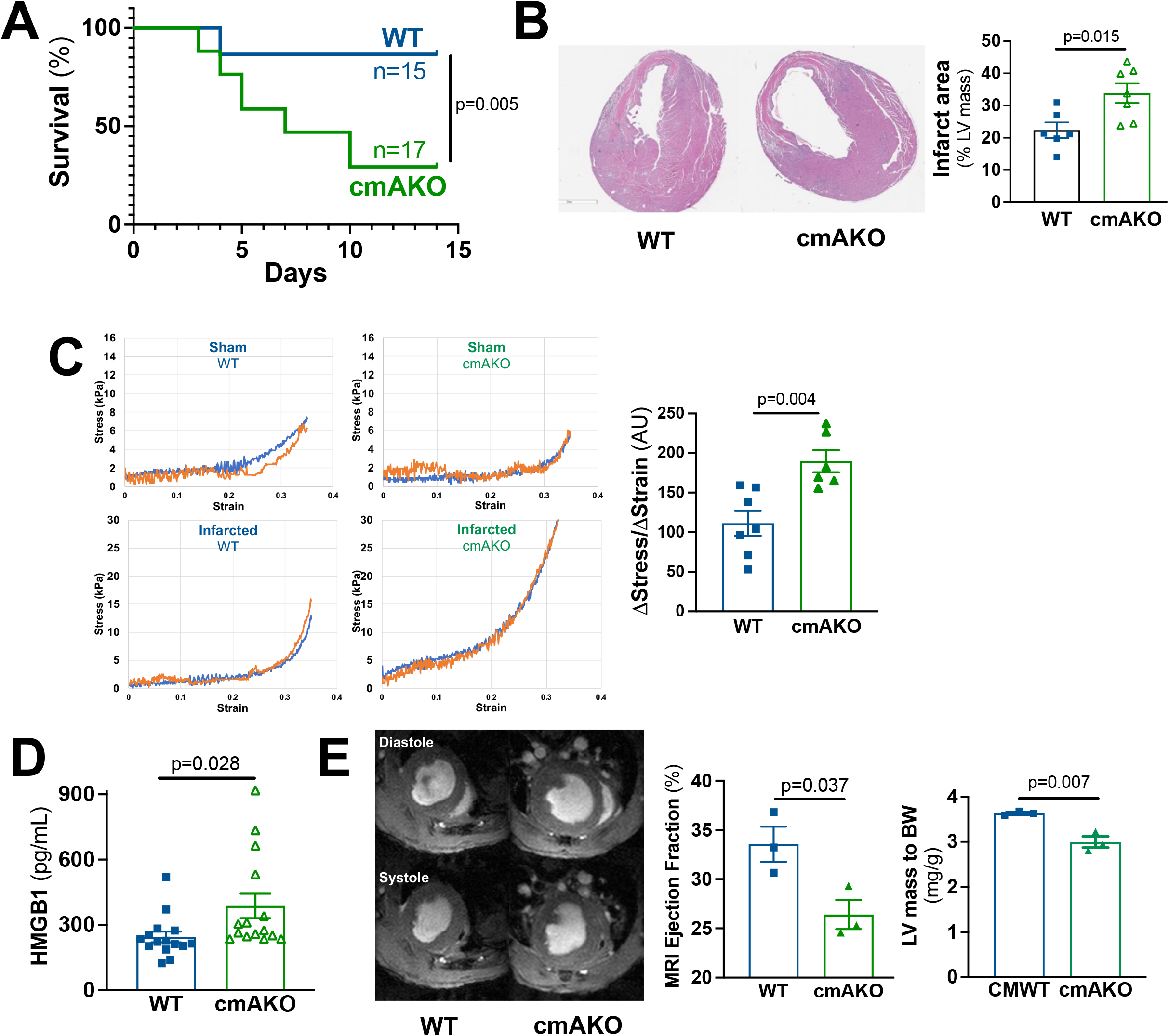
Disruption of the cardiomyocyte α1A-AR decreases survival and exacerbates cardiac dysfunction after experimental MI. WT and cmAKO mice underwent permanent ligation of the left coronary artery to create a myocardial infarction. **(A)** Kaplan-Meyer survival plot; **(B)** Infarct area calculated from summed slices in Image J; **(C)** Left ventricular mechanical properties as measured by a Biotester-5000 biaxial mechanical test apparatus; **(D)** ELISA compared serum abundance of high mobility group box 1 (HMGB1) in WT and cmAKO mice after infarction; **(E)** Cardiac MRI quantified left ventricular (LV) contractile function and structure, as indicated by ejection fraction and LV mass indexed to body weight (BW).

Changes in ventricular muscle tensile strength precede the onset of ventricular rupture in experimental acute MI(23). We employed a Biotester-5000 biaxial mechanical test system to determine whether mechanical properties differed in WT and cmAKO ventricles 3 days after myocardial infarction, prior to the onset of cardiac rupture. To validate this system, we initially compared infarcted myocardium with myocardium from sham-operated animals and found that infarcted tissue demonstrated an exponential stress-strain curve, consistent with previous reports.(24) We then compared the mechanical properties of sham-operated and infarcted WT and cmAKO mouse ventricles and found that the estimated biaxial tensile stretch (as in (25)) was >70% higher in cmAKO mice (190 ± 34 vs. 111 ± 42 kPa/% strain changes, p=0.0038, **Figure 2C and Supplemental Figure S2**). These data suggested that the absence of cardiomyocyte α1A-ARs compromises the biomechanical adaption of the left ventricle to experimental MI, potentially rendering the ventricle more susceptible to cardiac rupture(26).

We then sought to determine whether this significant mechanical alteration at Day 3 post-infarction was accompanied by other physiological indicators of injury. To determine whether infarct size correlated with circulating markers of cardiomyocyte death, we assayed serum high-mobility group box 1 (HMGB1). HMGB1 is released by cells undergoing necrotic cell death and is recognized as a sensitive biomarker that correlates to the extent of cardiomyocyte death in patients with MI(27). On Day 3 after LCA ligation, serum HMGB1 from cmAKO mice was 58% higher than in WT controls (387 ± 56 vs. 244 ± 25 pg/mL, p=0.028, **Figure 2D**). Using a small animal 9 Tesla magnet for cardiac MRI, we then compared cardiac morphology and contractile function of WT and cmAKO mice 3 days after LCA ligation. Contractile function in cmAKO mice was significantly worse than in WT after infarction (26 ± 3% vs. 34 ± 3%, p=0.037, (**Figure 2E)** with evidence of marked dyskinesis of the infarcted area **(Supplemental Video**). LV mass indexed to body weight was 18 ± 3% lower in cmAKO than WT mice, consistent with greater cell death (**Figure 2E**).

Collectively these findings demonstrate that the absence of cardiomyocyte α1A-ARs is associated with a pronounced susceptibility to myocardial injury and death after experimental myocardial infarction.

### Deficiency of cardiomyocyte α1A-ARs results in activation of RIP kinases and exaggerated cell death

Our histopathological (**Figure 2B**) and serum biomarker (**Figure 2D**) data indicated that infarct-associated necrotic cell death was exacerbated by the absence of the cardiomyocyte α1A-AR. Though cellular necrosis historically has been regarded as a passive process, emerging evidence has demonstrated clearly that programmed necrosis, or necroptosis, plays a critical role in numerous disease states including MI(28, 29). Necroptosis is initiated by binding of a ligand, often but not always a member of the TNFα superfamily, to a cardiomyocyte death receptor. Death receptor activation promotes the assembly of the necrosome that includes the receptor-interacting protein (RIP) kinases RIP1 and RIP3 and the terminal effector mixed lineage kinase domain-like protein (MLKL) leading to translocation of M LKL to the plasma membrane and necrosis. This signaling machinery is distinct from apoptotic signaling and activation typically requires intrinsic inhibition of apoptosis (reviewed in (28)).

To probe the contributions of necroptosis and apoptosis to the dramatic cmAKO phenotype, and to define whether cardiomyocyte α1A-ARs differentially constrained necroptosis and apoptosis in the setting of MI, we immunoblotted for canonical mediators of each process in WT and cmAKO mice 3 days after LCA ligation. RIP1 and RIP3 expression were 2-fold higher in the border zone of infarcted cmAKO hearts compared to infarcted WT controls (**Figure 3A**), consistent with activation of the necrosome. In contrast, the expression level of cleaved caspase-3 (c-Casp-3) and cleaved PARP was no different between the 2 groups (**Figure 3A**). We then asked whether the terminal necroptosis effector MLKL was differentially upregulated in the infarct border zone of cmAKO mouse hearts. Immunofluorescent staining revealed that MLKL was almost entirely absent in the myocardium of sham-operated control animals (**Figure 3B**). In contrast, the border zone of infarcted animals contained markedly higher expression of MLKL within cardiomyocytes, as defined using wheat germ agglutinin (WGA). Qualitative evaluation suggested that cmAKO hearts contained more MLKL than WT (Myh6-Cre^neg^/Adra1a^fl/fl^). Nuclear staining with DAPI identified robust infiltration of leukocytes consistent with the immune response that is activated by necrotic loss of cardiomyocyte membrane integrity. We then immunoblotted for MLKL to quantify its abundance in the border zone. We found that MLKL, relative to total protein, increased almost 8-fold in cmAKO animals but was not significantly changed in WT animals (**Figure 3C**).

**Figure 3.**
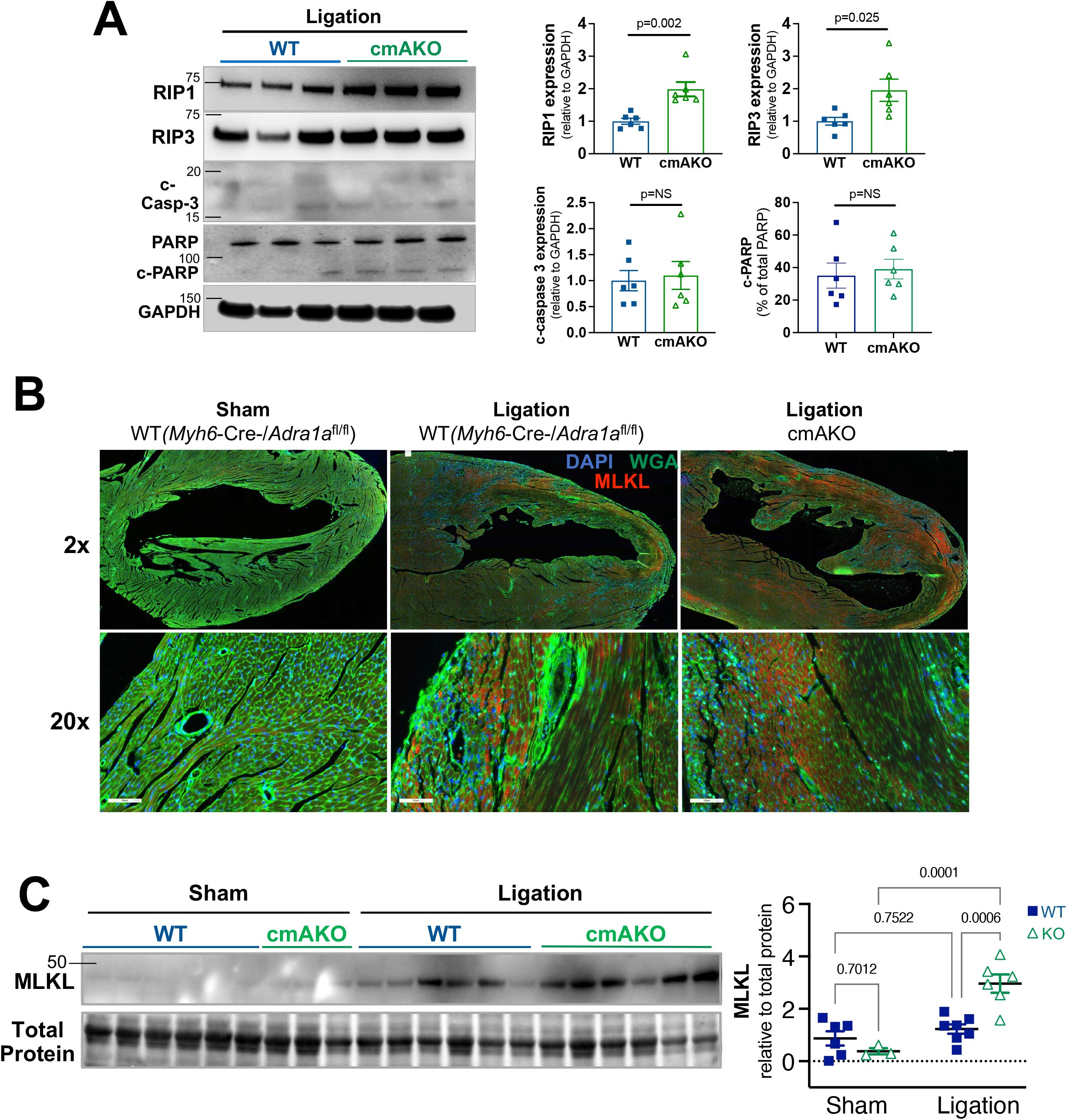
Mediators of necroptosis, but not apoptosis, are upregulated to a greater extent in cmAKO than WT mice after MI. Tissue was excised from the border zone of the left ventricular infarct 3 days after left coronary artery ligation. **(A)** Immunoblotting for mediators of necroptosis (RIP1 and RIP3) and apoptosis (c-PARP and c-Casp3) with representative image and summary densitometry; **(B)** Immunofluorescence for the necroptosis effector MLKL; **(C)** Immunoblotting for MLKL with representative images and summary densitometry.

Taken together, these results suggest that RIP1/3 mediated cardiomyocyte necroptosis, but not apoptosis, was exaggerated in mice lacking cardiomyocyte α1A-ARs after experimental MI.

### A selective α1A-AR agonist protects against RIP kinase-mediated cardiomyocyte death *in vitro*

To expand upon our *in vivo* findings demonstrating that the absence of cardiomyocyte α1A-ARs was associated with enhanced cell death and RIP kinase upregulation, we employed a well-established *in vitro* model of ischemia(30) induced by exposing primary neonatal rat ventricular myocytes (NRVMs) to oxygen and glucose deprivation (OGD). As indicated in **Figure 4A**, 24 hours of OGD resulted in significant cell injury as determined by LDH release when compared to standard culture conditions (34 ± 5 vs. 3 ± 1% of positive control, p<0.0001). Cell death was mitigated by selective α1A activation (A61603, 100nM) or by non-selective α1-AR activation with the combination of norepinephrine (NE, 100nM) and the beta-AR antagonist (β-blocker) propranolol (1μM). Next, we examined whether α1A-AR activation would protect against mitochondrial injury, as RIP3 activation promotes mitochondrial permeability transition pore (mPTP) opening and resultant loss of mitochondrial membrane potential (ΔΨ_m_)(31). Using the mitochondrial membrane permeant dye JC-1 (red aggregate with intact ΔΨ_m_, green monomer with compromised ΔΨ_m_) we found that 6 hours of OGD exposure led to a profound loss of ΔΨ_m_ that was partially rescued by coadministration of A61603 (205 ± 44% vs. 107 ± 37% aggregate/monomer, p=0.005, **Figure 4B**).

**Figure 4.**
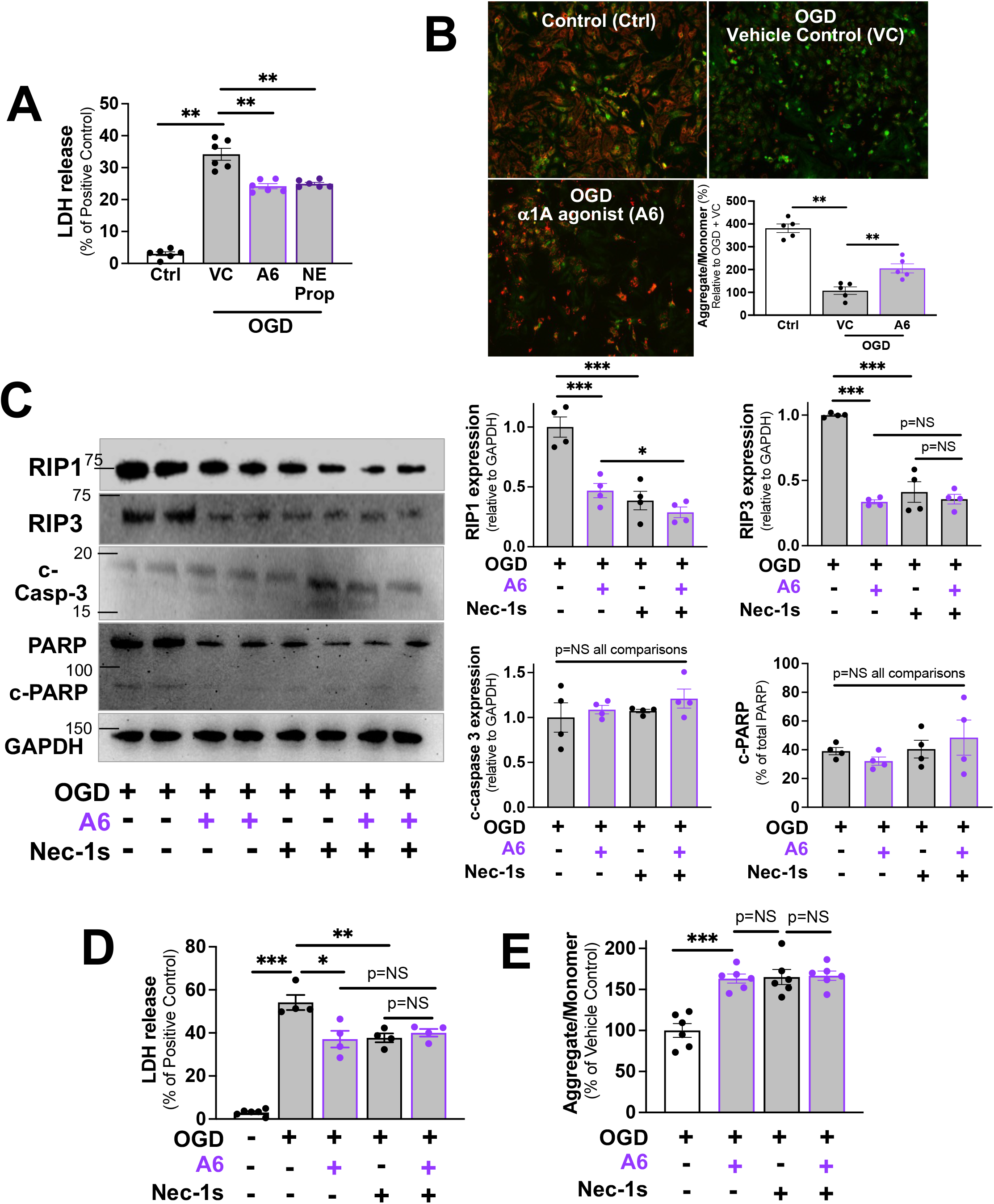
Pharmacologic activation of α1A-ARs protects neonatal rat ventricular myocytes from experimental ischemia. Ischemia was simulated in NRVMs using oxygen-glucose deprivation (OGD) for 24 hours in vehicle control (VC) with or without the α1A-AR agonist A61603 100nM or the non-selective AR agonist norepinephrine (NE, 1μM) with the β-AR antagonist propranolol (Prop, 1μM), or the RIP1 kinase antagonist Nec-1s 50μM. **(A)** Lactate dehydrogenase (LDH) measured cell death relative to positive control; **(B)** The mitochondrial membrane permeant dye JC-1 measured mitochondrial membrane potential (ΔΨ_m_) where red = intact ΔΨ_m_ and green = compromised ΔΨ_m_; **(C)** Cell lysates from OGD-exposed NRVMs in the presence or absence of A61603 or Nec-1s for 6h were immunoblotted. Image J measured relative densitometry; **(D)** LDH release measured OGD-exposed NRVM death after 6h in the presence or absence of A61603 or Nec-1s; **(E)** JC-1 measured NRVM ΔΨ_m_ in the presence or absence of A61603 or Nec-1s for 6h. *p<0.05, **p<0.01, ***p<0.001 by ANOVA with Tukey post-test.

To further define the importance of RIP kinases and necroptosis in our model systems, we used necrostatin-1stable (Nec-1s), a highly selective inhibitor of RIPK1(32). RIPK1 inhibitors, including Nec-1s have been shown to limit necrotic cell death due to ischemia in the heart and other organs(33, 34) and have been proposed as novel therapies for multiple disease processes mediated by ischemia(35). Here we exposed NRVMs to OGD in the presence or absence of the α1A agonist A61603 (A6) or Nec-1s 50μM (**Figures 4C-E**). A61603 and Nec-1s both decreased OGD-induced RIP1 by 50% and combination treatment further blunted RIP1 expression. Both A61603 and Nec-1s decreased RIP3 expression by 60-70%, but no additive effect was observed. Neither α1A activation nor RIP1 inhibition affected expression of the apoptosis mediators cleaved caspase-3 or cleaved PARP (**Figure 4C**). A61603 and Nec-1s both independently mitigated NRVM death (**Figure 4D**) and protected NRVM AΨ_m_ (**Figure 4E**) but no additive effect of α1A activation and RIPK1 inhibition was observed for these endpoints.

Collectively, these findings demonstrate that pharmacologic activation of the α1A-AR protects against ischemia-induced RIP kinase activation and necroptotic cell death. The absence of synergistic benefit from concomitant α1A activation and RIP inhibition raises the possibility that abrogation of necroptosis may contribute significantly to α1A-mediated cytoprotection. These *in vitro* results using NRVMs exposed to OGD are consistent with the findings from our LCA ligation model, suggesting that *in vivo* RIP kinase pathway activation likely results from cardiomyocyte necroptosis that is enhanced in the absence of the α1A-AR.

### Treatment with the RIP1 kinase inhibitor Nec-1s mitigates injury in cmAKO mice after LCA ligation

Our *in vitro* studies using NRVMs suggested that mitigation of necroptosis contributes to the cytoprotective effects of an α1A-AR agonist in the setting of ischemia. As an extension of these findings, we then sought to test whether inhibition of necroptosis *in vivo* could blunt the exaggerated response to injury in cmAKO mice using our experimental MI model.

To address this question, WT and cmAKO mice were administered Nec-1s (1.65 mg/kg body weight) or vehicle intravenously (IV) 10 mins prior to LCA ligation, followed by daily IV injection for two days prior to sacrifice on Day 3. Infarct size on Day 3 was no different in the cmAKO (20 ± 2% LV surface area) than the WT mice (16 ± 2%, **Figure 5A,** representative images **Supplemental Figure S3**), a contrast to the significant difference between genotypes in the absence of Nec-1s (WT 34 ± 8% vs. cmAKO 22 ± 6%, p=0.015, **Figure 2B**). Serum levels of the necrotic cell death marker HMGB1 were markedly higher in cmAKO than WT mice after infarct, but Nec-1s treatment abrogated that difference (**Figure 5B**). These findings suggested that necroptotic cell death accounted for the difference in HMGB1 release in WT and cmAKO mice and that HMGB1 may be a reliable biomarker for necroptosis post-MI. Consistent with findings from our previous experiments (**Figure 3A**), RIP1 (1.6-fold) and RIP3 (2-fold) were activated to a greater extent in the infarct border zones of cmAKO than WT mice (**Figure 5C**). Nec-1s treatment blunted RIP1 and RIP3 in all mice and abrogated the difference between the genotypes (**Figure 5C**).

**Figure 5.**
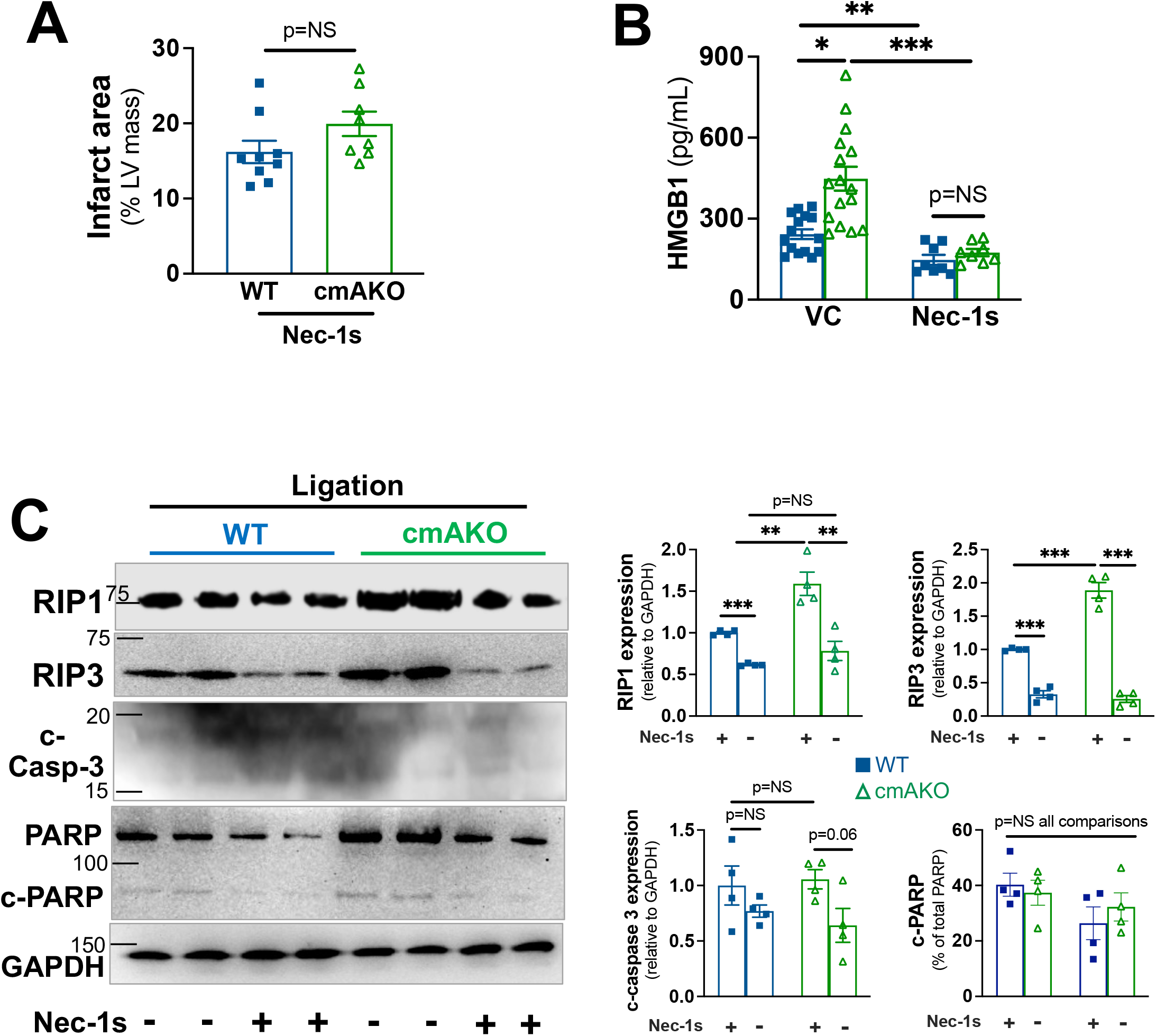
Treatment with the RIP1 kinase inhibitor Nec-1s mitigates the exaggerated response to injury in cmAKO mice subjected to left coronary artery (LCA) ligation. WT and cmAKO mice were injected intravenously (IV) with Nec-1s (1.65 mg/kg body weight) or vehicle 10 mins prior to LCA ligation, followed by daily IV injection for two days prior to sacrifice on Day 3. **(A)** Infarct area was calculated histologically; **(B)** Serum level of high mobility group box 1 (HMGB1), a measure of necrotic cell death was measured using an ELISA; **(C)** Lysates were prepared from the infarct border zones of all mice and immunoblotted for mediators of necroptosis (RIP1 and RIP3 kinases) and apoptosis (cleaved caspase-3 and cleaved-PARP). Summary densitometry was calculated in Image J. *p<0.05, **p<0.01, ***p<0.001 by ANOVA with Tukey post-test.

Taken together, these findings suggest that RIP1 kinase-mediated necroptosis contributes significantly to the enhanced susceptibility to ischemic injury in the absence of cardiomyocyte α1A-ARs.

### Alpha blocker exposure increases mortality after myocardial infarction in humans

Next we sought to determine whether the loss of α1-AR function also confers worse post-infarct outcomes in humans. Antagonists of α1-ARs (α-blockers) were developed to treat hypertension but currently are used most frequently to treat lower urinary tract symptoms due to benign prostatic hyperplasia in older men. Previous studies have identified an association between α-blocker use and incident HF(36, 37), but the effect of α-blockers on outcomes after MI has never been studied. We conducted a retrospective observational study using clinical data from the UNC Hospitals Percutaneous Coronary Intervention (PCI) Registry within the Carolina Data Warehouse for Health. We initially identified all patients (n=12,055) who were admitted to the UNC Healthcare system with a primary diagnosis of acute MI (ICD-9: 410 or ICD-10: I21) and underwent PCI between January 2014 and December 2018 (**Figure 6A**). From this population we merged instances with identical medical record numbers (the same patient admitted to the hospital multiple times) and filtered out patients younger than 60 years, as they rarely are prescribed α-blockers. The pre-specified primary outcome was defined as all-cause mortality prior to the end of the follow-up period (February 1,2020).

**Figure 6.**
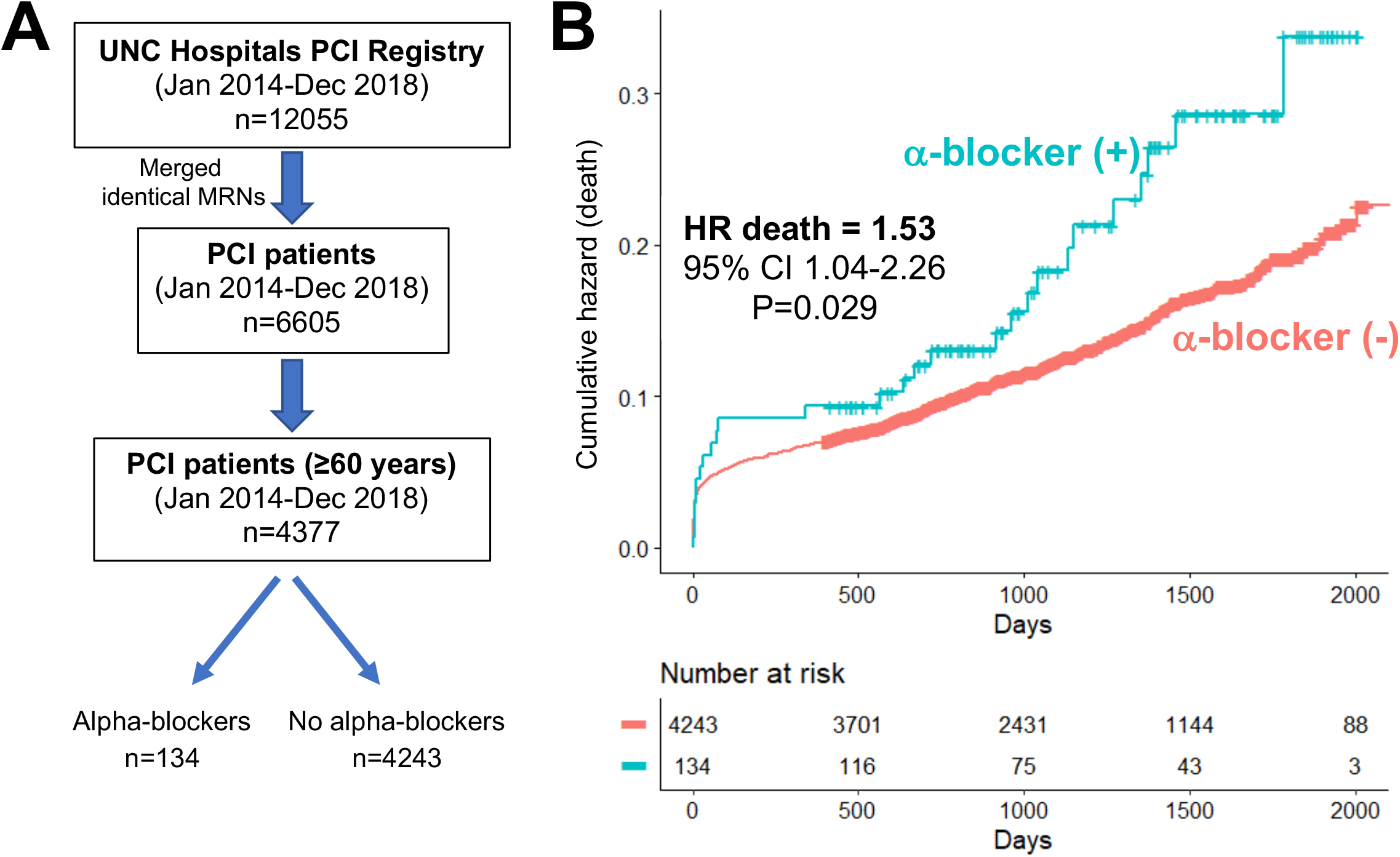
Use of α-blockers increased all-cause mortality among patients who were admitted to UNC Hospitals for acute myocardial infarction. **(A)** Patient selection algorithm for the retrospective observational study; **(B)** Kaplan-Meier survival analysis with hazard ratio (HR) for death, adjusted for age, sex, hypertension and heart failure.

This selection algorithm identified 134 patients taking α-blockers and 4243 patients not taking α-blockers at the time of their MI. The α-blocker (+) patients were older and more commonly male than the α-blocker (-) patients, but otherwise were clinically and demographically similar (**Table 2**). Our analysis found that the all-cause mortality rate was higher in α-blocker (+) patients (20.1% vs. 12.8%, p=0.019) during a median of 3 years of post-MI follow-up. Applying a Cox proportional hazards regression model with adjustment for age and sex, history of hypertension and history of heart failure, we found that the hazard ratio for death in the α-blocker (+) post-MI patients was 1.53 (95% CI: 1.04-2.26, p=0.029) (**Figure 6B**).

**Table 2.**
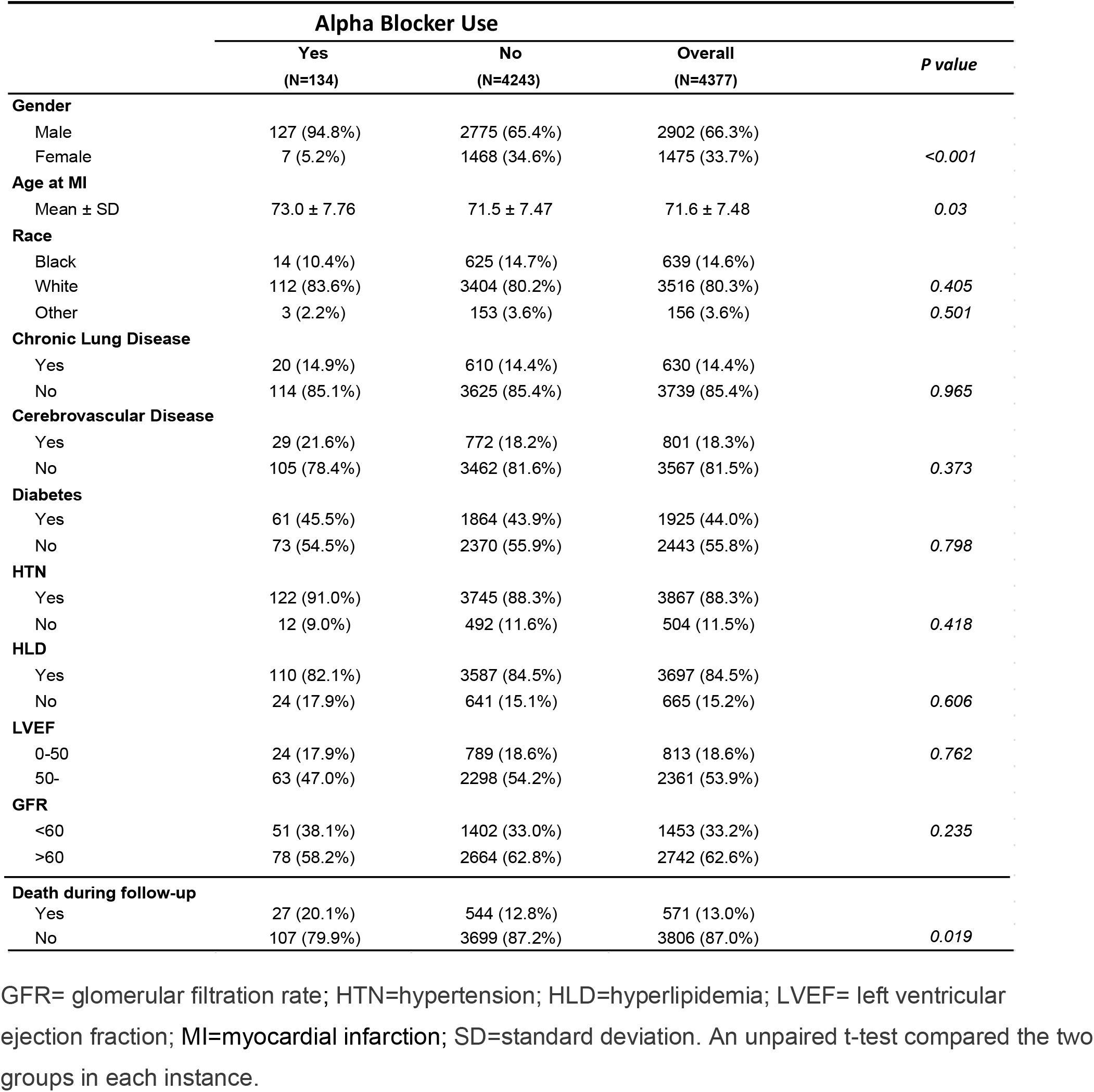
Clinical and demographic characteristics of patients ≥ 60 years old who underwent percutaneous coronary intervention after admission to UNC Hospitals for acute myocardial infarction.

## Discussion

In this study we show that mice that lack the α1A-AR subtype only in cardiomyocytes are at a significantly elevated risk of death within the first 5 days after experimental MI. These new cmAKO mice experienced larger infarcts and exaggerated ventricular remodeling and contractile dysfunction within 3 days post-infarct. *In vitro* studies corroborated these findings, demonstrating that α1A-AR activation limits cell death in the setting of ischemic injury. Both *in vivo* and *in vitro* studies suggested that constraint of necroptosis contributes to cardioprotection by α1A-ARs. A retrospective observational clinical study at our institution indicated that the use of medications that block α1-ARs was associated with increased risk of mortality after MI, offering a potential clinical parallel for our experimental findings.

To the best of our knowledge, our study is the first to demonstrate an essential protective role for endogenous cardiomyocyte α1A-ARs in response to injury using a cardiomyocyte-specific loss-of-function model. Mice with 66-fold overexpression of rat *Adra1a* have improved survival and contractile function 15 weeks after LCA ligation(17). Transgenic overexpression (15-40 fold) of the α1A-AR in rat cardiomyocytes confers enhanced ischemic second window ischemic preconditioning(38) and is associated with decreased ventricular remodeling and enhanced angiogenesis 4-6 weeks after MI(18). Global α1A-AR knockout mice have increased mortality and enhanced ventricular remodeling 4 weeks after MI(19). Here we have created a cardiomyocyte-specific α1A-AR knockout mouse line to avoid the potentially confounding effects of either supraphysiological protein abundance in the heart or systemic effects from global deletion of a widely expressed receptor. Our cmAKO mice have normal basal cardiac structure and function (**Figure 1**, **Table 1**), similar to both the global α1A-KO mice(19) and to a recently published line with inducible cardiomyocyte-specific α1A-AR deletion(39). Though deletion of cardiomyocyte α1A-ARs does not alter basal phenotype appreciably, it does confer significantly enhanced susceptibility to injury, as evidenced by 50% mortality within 5 days of MI.

Here we have by necessity focused our mechanistic analyses on an earlier time point given the excessive early mortality due primarily to cardiac rupture (66.7%, 6/9) as confirmed by standardized methodology(21). Our observations from biaxial tensile tests confirmed that disruption of cardiomyocyte α1A-ARs leads to maladaptive biomechanical properties predisposing to elevated intracardial pressure and free wall rupture. Though the propensity for post-MI cardiac rupture likely is multifactorial, extensive necrotic loss of cardiomyocytes predisposes myocardium to rupture via proinflammatory responses and dynamic changes in the extracellular matrix (40–42). Historically, necrosis has been considered an “unregulated” and passive form of cell death. However, studies carried out over the past two decades have unequivocally characterized necroptosis as a regulated form of cellular necrosis(43) that plays critical roles in multiple types of cardiac pathology(28), including myocardial ischemia and infarction(31, 44, 45).

The peri-infarct period is characterized by elevated levels of endogenous catecholamines(46) that activate ARs on cardiomyocytes. High concentrations of the β-AR agonist isoproterenol (85 mg/kg s.c.) induce cardiomyocyte necrosis with induction of RIP1 and RIP3 in mice(47), suggesting that β-AR hyperactivation may promote necroptosis to contribute to the pathobiology of MI. Regulation of necroptosis by α1-ARs has not been studied previously. Here, we report that endogenous α1A-ARs protect cardiomyocytes after MI in part by constraining RIP kinase-MLKL-mediated necroptosis, but not apoptosis. Our findings would appear to be inconsistent with a previously published study showing enhanced myocardial apoptosis after experimental MI in global α1A-AR KO mice(19). However, there are several important differences in the design of these studies that may explain the apparent discrepancies. Yeh et al analyzed the infarcted heart 4 weeks after LCA ligation whereas here we carried out most of our studies in mice 3 days after ligation. This temporal difference may be important given that the predominant modes of cell death vary with time after infarction(48). Secondly, Yeh et al assayed cell death from remote non-infarcted myocardium, whereas we exclusively utilized tissues from peri-infarct border zone. Lastly, it is entirely plausible that their findings were influenced by systemic loss of α1A-ARs, whereas we employed a cardiomyocyte-specific KO mouse model.

Our study also is the first to report on enhanced risk of mortality in patients taking α-blockers, though other studies have identified an association between α-blockers and incident heart failure. In the landmark ALLHAT (Antihypertensive and Lipid Lowering treatment to prevent Heart Attack Trial), subjects taking doxazosin had a two-fold higher risk of developing heart failure than patients taking chlorthalidone(36). Partly as a result of this finding, non-selective α-blockers are no longer considered first-line therapy for hypertension. However, α-blockers remain the most prescribed medication for managing symptomatic benign prostatic hyperplasia (BPH), which affects over 70% of US males greater than 60 years old(49). In fact, the Medical Expenditure Panel Survey (Agency for Health Care Research and Quality) estimates that over 5 million patients were prescribed the α1A-AR selective antagonist tamsulosin in the United States in 2019. A recent population-based study from Canada found that men taking α-blockers for BPH had an increased risk of developing heart failure (HR 1.22, 95% CI 1.18-1.26) when compared to men with BPH who were not on medical therapy(37). Neither of the previous studies reported increased mortality related to α-blockers as we do, though we studied a particularly high-risk population in post-MI patients.

Our study has several important limitations. The complex pathobiology of myocardial infarction is both spatially and temporally heterogeneous. We chose by necessity to focus our analyses on selected post-infarct timepoints and were painstaking in our efforts to study infarct border zone, but we cannot exclude the possibility that the α1A-AR regulates distinct processes at other times or in different myocardial locations after infarct. Indeed, it is quite possible that alternative mechanisms other than necroptosis may contribute to the cardioprotective effects of cardiomyocyte α1A-ARs. For example, rats with cardiomyocyte-specific α1A-AR overexpression exhibited attenuated post-infarct remodeling characterized in part by enhanced formation of coronary collaterals mediated by vascular endothelial growth factor A(18). We also cannot exclude the possibility that developmental effects contributed to the exaggerated susceptibility to injury in our cmAKO mice. In this regard, it would be interesting to study the response in a conditional knock down model such as the one created by Graham and colleagues recently(39). A conditional model knockout model would have better fidelity to human α-blocker exposure and would allow for explication of the role of α1A-ARs in the more remote post-infarct period. Finally, we acknowledge that our clinical study is relatively small and drawn from a single medical center. It clearly will be important to determine whether the increased risk of post-MI mortality can be replicated in larger datasets of patients taking α-blockers.

Despite these limitations, our study significantly advances our understanding of the mechanisms underlying the cardioprotective effects of the α1A-AR, identifying a previously unrecognized connection with necroptosis. The marked susceptibility to injury of our novel cardiomyocyte-specific α1A-AR knockout mouse suggests that enhanced cardiovascular risk in patients taking α-blockers may arise directly from inhibition of adaptive functions of cardiomyocyte α1A-ARs rather than systemic effects.

## Author contributions

**Research study design:** JDZ, JSR, JCS, BCJ

**Conducting experiments:** JDZ, PBS, WH, LO, AJS, TA, HY, BEM

**Acquiring data:** JDZ, PBS, WH, LO, AJS, TA, HY, BEM

**Analyzing data:** JDZ, SHL, YYS, HSH, JSR, JCS, BCJ

**Providing reagents:** PCS, BEM, JCS

**Manuscript preparation:** JDZ, BCJ, PCS

## Methods

### Mice

C57BL/6J and ROSA^mT/mG^ (stock # 007676) mice were purchased from Jackson lab (Bar Harbor, Maine). The cmAKO mouse line was generated by breeding *Myh6*-Cre (originally from the lab of E. Dale Abel at the University of Iowa(20) and provided by Leslie Leinwand at the University of Colorado(50)) to *Adra1a* flox/flox mice with loxP sites flanking the first coding exon (constructed in the Paul C. Simpson lab at University of California, San Francisco/San Francisco VA Medical Center). All mice were backcrossed regularly and maintained on a C57BL/6 genetic background. 12–16-week-old males were used in all experiments. Floxed mice (αMHC-Cre^neg^/α1A^fl/fl^) were used as WT controls for cmAKO mice.

### Quantitative reverse transcriptase PCR (qRT-PCR)

Total RNA was isolated from cells and tissue (Qiagen RNeasy Plus mini kit #74134) and analyzed using a NanoDrop (ThermoScientific). For qRT-PCR, one μg of RNA was reverse transcribed using High Capacity cDNA Reverse Transcription Kit (Life Technologies #4368814). Two step qRT-PCR reactions contained 2% of the cDNA product. All reactions were performed in triplicate in a Roche 480 Light Cycler. Relative quantitation of PCR products used the ΔΔCt method relative to two validated reference genes (*Tbp* and *Polr2a*). Similar efficiencies were confirmed for all primers. All probes and primers were from Roche. Primer sequences:

Tbp F:ggcggtttggctaggttt; R:gggttatcttcacacaccatga; UPL Probe # 107
Polr2a F:aatccgcatcatgaacagtg, R:tcatcatccattttatccacca; UPL Probe # 69
Adra1a F: attgtggtgggatgcttcgtcct; R: tgtttccggtggcttgaaattcgg; UPL Probe # 105
Adra1b F: ttcttcatcgctctcccact; R: gggttgaggcagctgttg; UPL Probe # 20

### Echocardiography

Transthoracic echocardiography was performed on conscious, loosely restrained mice in the McAllister Heart Institute Rodent Surgical Core using a VisualSonics Vevo 2100 ultrasound system (VisualSonics, Inc., Toronto, Ontario, Canada). Two-dimensional and M-mode echocardiography were performed in the parasternal long-axis view at the level of the papillary muscle. Left ventricular systolic function was assessed by fractional shortening (%FS = [(LVEDD -LVESD)/LVEDD] × 100). Reported values represent the average of at least five cardiac cycles per mouse. Sonographers and investigators were blinded to mouse treatment condition during image acquisition and analysis.

### Mouse myocardial infarction model

Mice were subjected to permanent left coronary artery (LCA) ligation as previously described (51) and subsequently assessed for left ventricular (LV) morphology and function. For LCA ligation as well as cardiectomy, mice were anaesthetized by inhalation of isoflurane (2%). For postoperative analgesia, 5mg meloxicam/kg body weight was applied every 24 hours for first 72 hours post-surgery. For the Nec-1s (7-Cl-O-Nec-1, Sigma) studies, vehicle or Nec-1s (1.65 mg/kg body weight) was given intravenously (IV) 10 minutes prior to LCA ligation, followed by daily IV injections for 2 days.

### Immunostaining and histological analysis

To fix heart tissue for immunohistochemistry, mice were heparinized and the heart was perfused with 10 mL of PBS followed by 20 mL of 4% paraformaldehyde (PFA)-PBS through a 23-gauge butterfly needle, then excised and placed in 4% PFA-PBS for 24 h before transfer to 70% ethanol. Hearts were sectioned and stained using standard methods in the University of North Carolina Histology Research Core. Slides were scanned using an Aperio ScanScope (Aperio Technologies, Vista, CA) and analyzed in Aperio ImageScope software. For immunohistochemistry, hearts were fixed overnight in 4% PFA-PBS, incubated in 30% sucrose-PBS, and then frozen in OTC medium (Tissue-Tek, Hatfield, PA). Frozen sections (10 μm) obtained with a Leica cryostat (Leica, Buffalo Grove, IL) were placed on glass slides, dried at room temperature, and then incubated with primary antibodies (c-caspase-3, RIP kinase). After washing, sections were incubated for 3h at room temperature with secondary antibodies (1:1,000, Life Technologies, Grand Island, NY). Fluorescence was observed with a Zeiss LSM710 confocal microscope.

### Histopathological measurement of infarct size

Infarct size was measured as previously described(22) and 6-8 mouse hearts in each group were utilized for analysis. Briefly, each LV was sliced in the transverse axis from the apex to the base using a microtome at 5 μm thickness with an interval of 300 μm between each section. All sections were mounted on glass slides and stained with hematoxylin and eosin for further quantitative analysis. Slides were scanned using an Aperio ScanScope (Aperio Technologies, Vista, CA) and analyzed in Aperio ImageScope software. ImageJ software was used to measure areas of affected (infarction and infiltration) and whole LV myocardium. Affected area and the total area of LV myocardium were traced manually in the digital images by an investigator who was blinded to the genotype of the sample, then measured automatically by the computer. Infarct size, as expressed as a percentage of total LV myocardial surface area, was calculated by dividing the sum of injured areas from all sections by the sum of LV areas from all sections (including those without infarct) and multiplying by 100%.

### Cardiac magnetic resonance imaging (MRI)

Anesthesia was induced by using 3% isoflurane followed by ≤ 2% isoflurane for maintenance as required. Respiration rate was used to monitor depth of anesthesia and was maintained between 20-70 breaths per minute. Body temperature was monitored and controlled within the range of 37±2°C. Cardiac MRI, gated to EKG, was obtained in prone position using a 9 Tesla magnet equipped with a 370 mT/m gradient system, a 38 mm birdcage quadrature coil, and ParaVision 6.0 software. To assess cardiac function, short-axis images were acquired with the retrospectively-gated FLASH sequence with the following settings: 7-9 slices; field-of-view: 4.5 x 4.5 x 0.1cm; image matrix size: 256÷256; repetitions: 100; flip angle: 15°; echo time: 2.05ms; repetition time: 8.91ms; number of reconstructed frames: 8. A single person contoured the ventricles, while a second person checked the quality of the contouring. In case of disagreement the contouring was adjusted to achieve mutual agreement. End-systolic (ESV) and end-diastolic (EDV) cardiac phases were automatically determined for quality check of the cardiac contouring. A maximum mass difference of the left ventricle of 10% was allowed between both cardiac phases. The heart rate (HR) was extracted from the IntraGate sequence. These parameters were used to calculate the stroke volume (SV = EDV -ESV), cardiac output (CO = SV x HR) and ejection fraction (EF = SV/EDV x 100). All measurements and calculations were made by investigators blinded to genotype.

### Mechanical testing of ventricular myocardium

Mechanical testing was performed using a Biotester-5000 biaxial test system (Cellscale, Waterloo, Canada) following a previously reported method (52). In brief, at day 3 post-MI, male mice were sacrificed. The left ventricle was separated from the right ventricle and atria, and then cut into half along the long axis, creating a “sheet” of ventricular tissue. The isolated tissue was mounted biaxially on the Biotester with the infarct centered(52), then stretched according to manufacturer’s protocols. Deformation was measured by image tracking analysis. Stresses and corresponding strains were fit to a four parameter Fung-type model where the strain energy density was calculated to yield the Cauchy stress. Tissue stiffness was quantified as the slope of the Cauchy stress-stretch ratio curve.

### Serum HMGB1 ELISA

Blood was obtained from the inferior vena cava at the time of sacrifice 3 days after LCA ligation. Detection of serum HMGB1 was performed using a commercially available kit (Novus Biologicals, Centennial CO).

### Immunoblotting

Tissue and cell lysates were produced in RIPA buffer supplemented with PhosSTOP (Roche Diagnostics, Indianapolis, IN) and protease inhibitor cocktail (Roche Diagnostics). Subsequently, samples were incubated in 4× LDS sample buffer, including 2% β-mercaptoethanol, for 10 min at 70°C. SDS-PAGE and immunoblot analysis were performed using the 4–12% Nupage gel system (Life Technology, Foster City, CA). Membranes were blocked in 5% milk-Tris-buffered saline-Tween 20 and incubated in primary antibody overnight at 4°C and then secondary horseradish peroxidase (HRP)-conjugated antibodies for 1 h at room temperature. Images were generated using Amersham ECL Select Western Blotting Detection Reagent (GE Healthcare Life Sciences, Marlborough, MA) and the ChemiDoc™ Imaging System (Biorad, Hercules, CA).

### Antibodies

PARP/cleaved PARP Cell Signaling 9542; CAMKIId Abcam #181052; pCAMKIId Abcam #ab32678; RIPK3 Cell Signaling #95702; pRIPK3 Abcam #209384; RIP1 Cell Signaling #3493s; pRIP1 Cell Signaling #31122; MLKL Millipore Sigma #MABC604; pMLKL Abcam #196436m

### Neonatal rat ventricular myocytes (NRVMs)

Female Sprague-Dawley rats and newborn litters were obtained from Charles River. NRVMs were isolated as previously described(53). Experiments were carried out after 36–72 h of serum starvation in the presence of insulin, transferrin, and BrdU.

### Lactate dehydrogenase (LDH) assay

The supernatant from cultured NRVMs was collected to determine the content of LDH via cytotoxicity detection kit (Roche, USA), following the manufacturer’s instructions.

### Mitochondrial membrane potential (ΔΨm) assay

Mitochondrial membrane potential in NRVMs was measured by 5, 5’, 6, 6’-tetrachloro-1, 1’, 3, 3’-tetraethylbenzimidazolylcarbocyanine iodide (JC-1) reduction. Cells were stained with JC-1 (Abcam, Cambridge, MA) according to the manufacturer’s protocol. In brief, after appropriate treatment, serum-starved NRVMs were exposed to JC-1 at 2 μM for 30 min. Cells were washed once with medium then analyzed by plate reader (CLARIO star, BMG LABTECH, Germany). JC-1 green fluorescence was excited at 488 nm and emission was detected using a 530 ± 40 nm filter. JC-1 red fluorescence was excited at 488 nm and emission was detected using a 613 ± 20 nm filter. A shift in fluorescence from red (JC-1 aggregates) to green (JC-1 monomers) indicates JC-1 dissociation from mitochondria into the cytosol, implying a reduction of ΔΨm.

### Retrospective cohort clinical study

We conducted a retrospective observational study using clinical data from the Carolina Data Warehouse for Health. The study population consisted of all patients who were admitted to the UNC Healthcare system with a primary diagnosis of acute myocardial infarction (ICD-9 code: 410 or ICD-10 code I21) and underwent percutaneous coronary intervention between January 2014 to December 2018. Patients were grouped according to whether they were discharged from the hospital with a prescription for an α-blocker. The pre-specified primary outcome was defined as all-cause mortality prior to the end of the follow-up period (February 1, 2020). A Cox proportional hazards regression model was used for statistical analysis. Descriptive statistics were used for baseline characteristics. Models were adjusted for demographic or clinical values that were significantly different between groups at baseline (age, sex, history of heart failure, history of hypertension). All statistical analysis was done using R (R-project).

### Statistics

The values of each parameter within a group are expressed as the mean ± SEM. Statistical analysis were performed by using ANOVA followed by Tukey post hoc tests (3 or more groups) or by Student t-tests (2 groups) using GraphPad Prism 9. A p-value < 0.05 was considered statically significant. **Study Approval:** Animal care and experimental protocols were approved by the University of North Carolina at Chapel Hill Institutional Animal Care and Use Committee and complied with *Guide for the Care and the use of Laboratory Animals* (National Research Council Committee for the Update of the Guide for the Care and Use of Laboratory Animals, 2011). The retrospective cohort clinical study was approved by the UNC Institutional Review Board (IRB 19-3324). Written informed consent was obtained from all subjects prior to data collection.

---

**Supplemental Figure S1.**
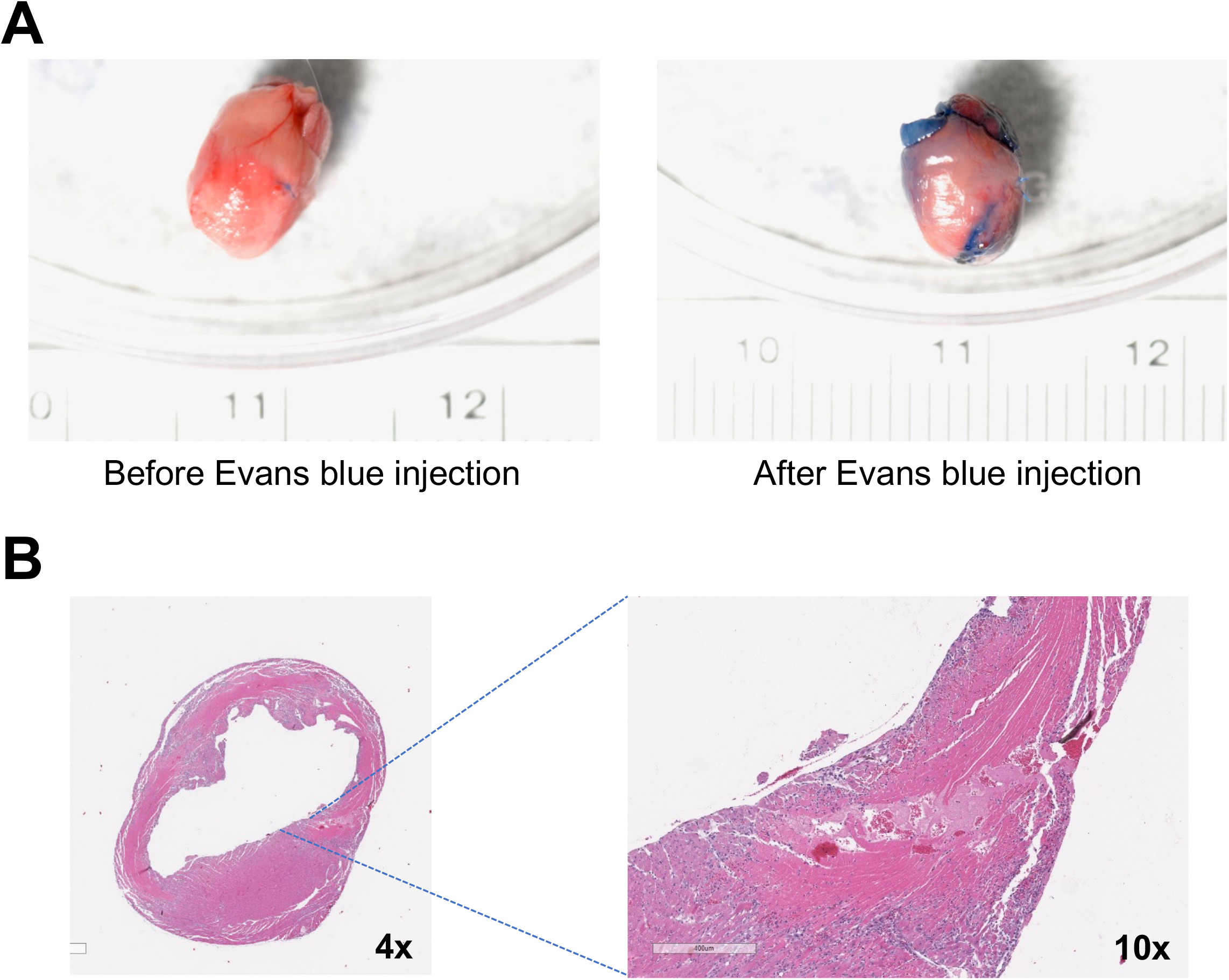
After infarction, 67% of cmAKO mice died of ventricular rupture. **(A)** Representative gross anatomical images; **(B)** Representative H&E images

**Supplemental Figure S2.**
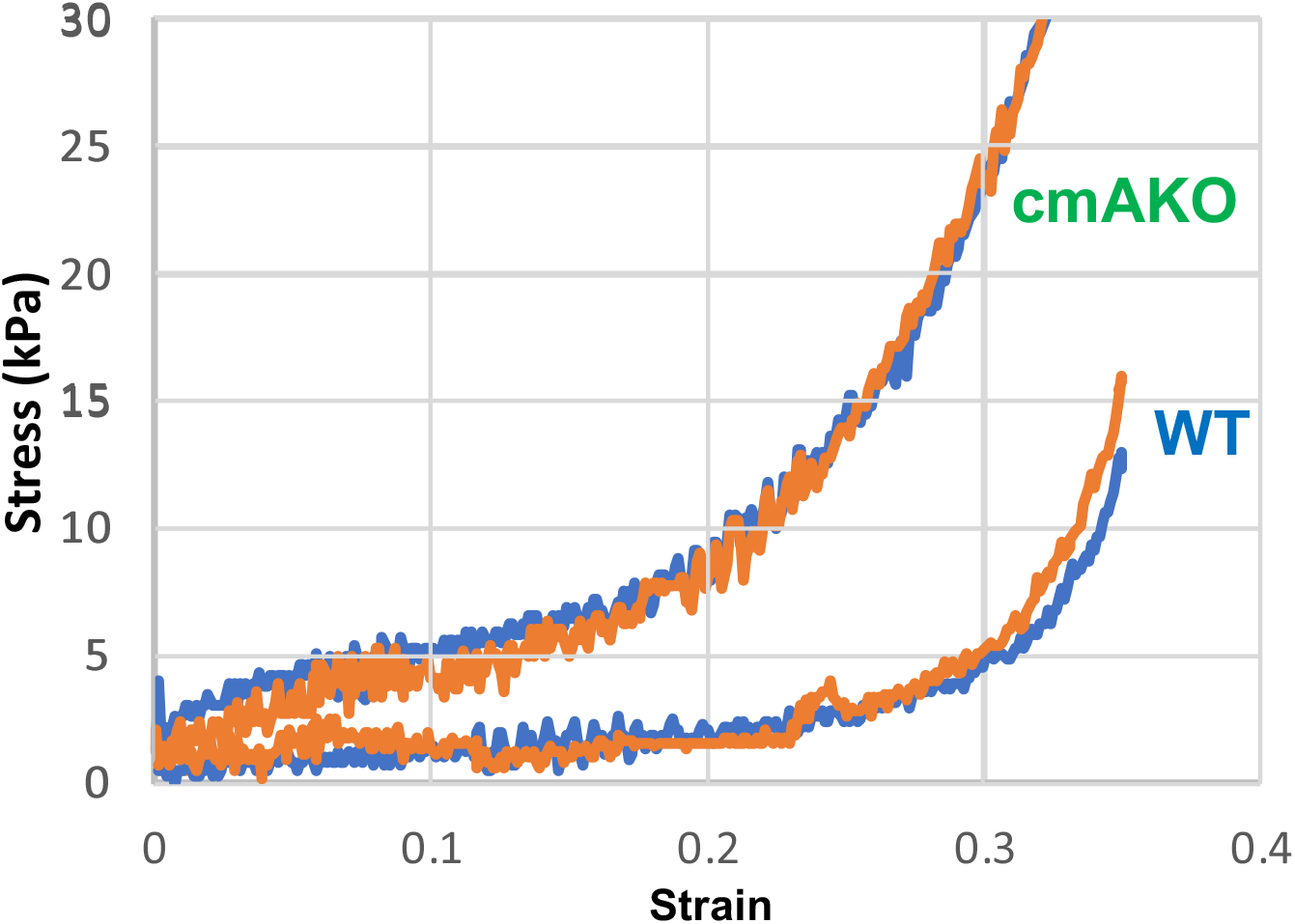
Representative plots of stress-strain curves for left ventricular myocardium 3 days after experimentally induced myocardial infarction. Orange—plots from X axis, blue—plots from Y axis.

**Supplemental Figure S3.**
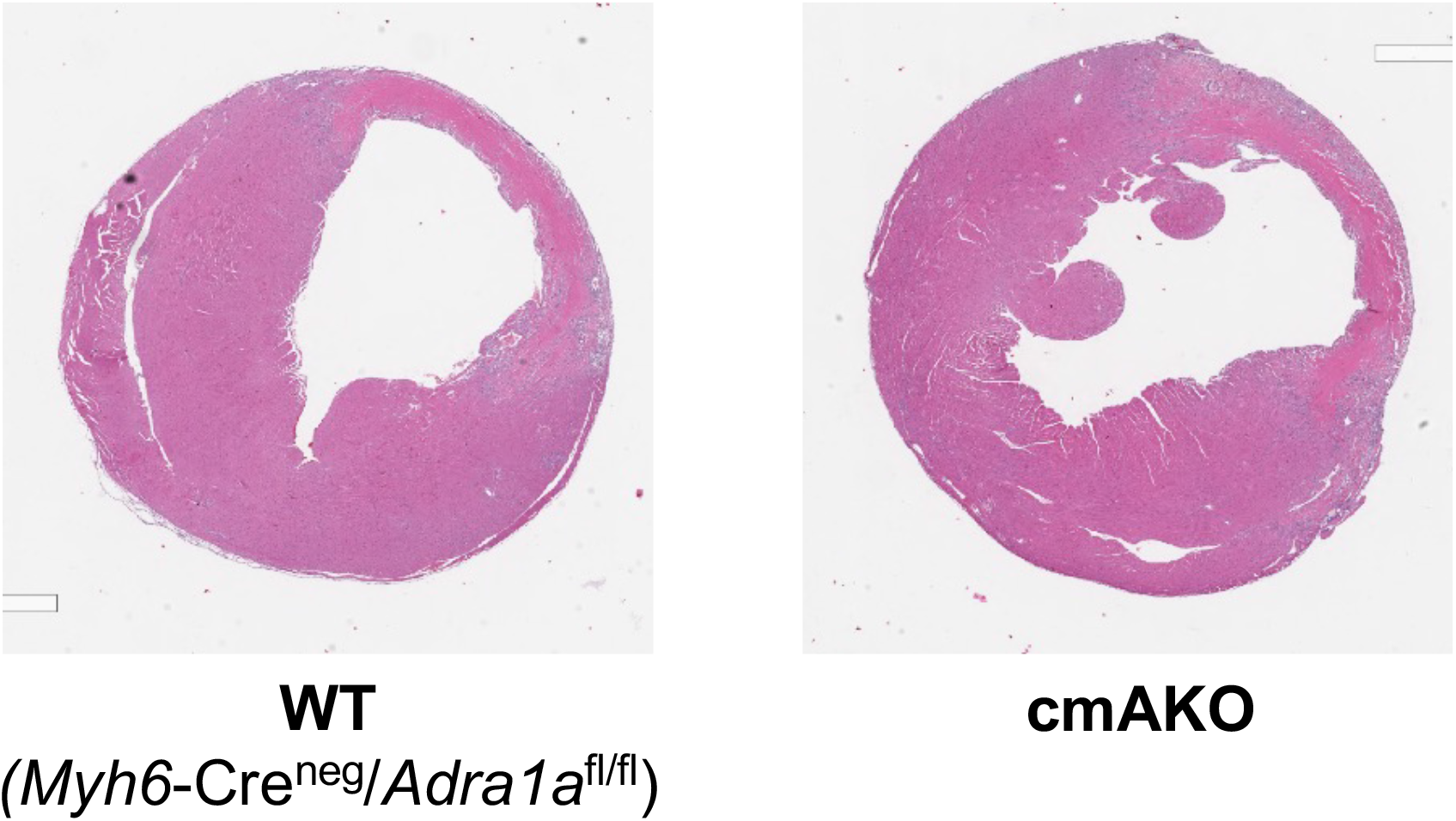
Representative H&E staining of post-infarct Day 3 hearts exposed to treatment with the RIP1 inhibitor Nec1s

